# Ecosystem connectivity and configuration can mediate instability at a distance in metaecosystems

**DOI:** 10.1101/2023.03.03.531041

**Authors:** Christina P. Tadiri, Jorge O. Negrín Dastis, Melania E. Cristescu, Andrew Gonzalez, Gregor F. Fussmann

## Abstract

Ecosystems are connected by flows of nutrients and organisms. Changes to connectivity and nutrient enrichment may destabilise ecosystem dynamics far from the nutrient source. We used gradostats to examine the effects of trophic connectivity (movement of consumers and producers) versus nutrient-only connectivity in different metaecosystem configurations (linear vs dendritic) on dynamics of *Daphnia pulex* (consumers) and an algae (resources). We found that peak population size and instability (coefficient of variation; CV) of *Daphnia* populations increased as distance from the nutrient input increased, but were lower in metaecosystems connected by all trophic levels compared to nutrient-only connected systems and in dendritic systems compared to linear ones. We examined the effects of trophic connectivity (i.e. how many trophic levels are moving rather than one or the other) using a generic model to qualitatively assess patterns of ecosystem dynamics. Our model predicted increased population sizes and fluctuations in ecosystems with nutrient connectivity, with this pattern being more pronounced in linear rather than dendritic systems. These results confirm that connectivity may propagate and even amplify instability over a metaecosystem to communities far from the source disturbance, and suggest a pathway for future experiments, that recreate conditions closer to those found in natural systems.

## Introduction

Ecosystem stability is a central theme in ecology with a large body of theory focused on identifying the key processes that stabilise versus those that destabilise an ecosystem’s dynamics (1-3). A significant body of research now invokes spatial processes, such as species dispersal and nutrient flows that may alter their movement in space over time, as key governors of stability in trophic systems (4). Anthropogenic perturbations to stability have been addressed and include altered climate gradients (5), altered trophic structure by over-exploitation (6), altered rates and scales of nutrient flows (7), and permanent shifts in the size (8) and connectivity of habitats (4). Experiments have contributed significantly to our understanding of the factors driving instability (9, 10), but few have sought to identify how these factors interact to define stability across trophically structured metaecosystems (11).

Altered patterns and rates of ecosystem connectivity can cause instability over large spatial extents both locally and at the metacommunity level. Here instability is any form of dynamics that threatens the persistence of a species locally or across the metacommunity. Connectivity via the spatial flows of nutrients, resources and consumers can carry or even amplify the destabilising effects of point sources of disturbance and perturbation over large distances (7, 12). For example, changes in the rates of movement of consumers may synchronise consumer dynamics driving instability (13), leading to greater metacommunity instability, as measured by increases in the variance in population dynamics within and among local communities (12).

Nutrient enrichment is another well-known source of community instability. This phenomenon, known as the paradox of enrichment (14) causes ecosystems to transition from stable dynamics to a regime characterised by large oscillations in the dynamics of primary producers and consumers. The consumer-resource oscillations caused by nutrient enrichment can increase the risk of extinction especially of edible species, allowing inedible species to become dominant (15). From this theory we expect connectivity and nutrient enrichment to interact strongly to define patterns of instability over entire metacommunities.

McCann et al (7) derived theoretical results in networked communities showing how ecosystem connectivity due to nutrient or consumer movement may allow nutrient enrichment at a point source to accumulate and drive instability at a terminal community far from the source. They also demonstrated the differential effects of either nutrient or consumer movement on system stability, where nutrient movement alone may be stabilising but lead to increases in consumer population sizes, whereas movement of consumers alone leads to amplification of consumer populations in the terminal community which drives instability. Additionally, they predicted that heterogeneity in the flow rates across a connected metaecosystem can be stabilising. They also highlighted empirical observational examples of nutrient-driven instability at a distance with nutrient movement, though none with consumer movement alone. These theoretical predictions have not been tested with a controlled experiment.

In this study we combined a replicated gradostat experiment and model simulations to test how metaecosystem configuration, heterogeneity, and functional connectivity alter the stability of the community dynamics. Modelled after the chemostat, a gradostat apparatus consists of a series of flasks connected by tubing through which medium is pumped in, through and out in order to create a sustained nutrient gradient (16). Gradostats thus offer the ability to experimentally test predictions about metaecosystems in microcosms; they have been used to examine competition dynamics (17-19) and consumer-resource dynamics (16, 20) in metacommunities. The direction and speed of flow and configuration of the flasks (‘nodes’) in the gradostats metaecosystem network may be modified to a greater degree than in chemostats. The gradostat offers high control over spatial flows in a lab setting and the small size allows for multiple treatments in a replicated experimental design.

To identify metaecosystem dynamics characteristic of our system we studied *Daphnia*-algae-nutrient interactions in gradostats with two metaecosystem configurations—linear and dendritic networks—and two levels of connectivity (only connected by flow of nutrients or connected by movement of all trophic levels). We hypothesised based on the findings of McCann et al (7) that instability would be greatest in the terminal node of gradostats, and that this effect would be stronger in those gradostats where consumers and resources move in addition to nutrients. We further hypothesised that metaecosystem configuration would modify instability in the terminal node, with those in dendritic configurations being more stable than linear ones due to the slower upstream nodes dampening the influx of nutrients and consumers to the terminal node. We compare our experimental results to simulations of a generic consumer-resource model describing the experimental design.

## Methods

### Laboratory Experimental Design

For a full description of culture and treatment preparation, see Supplemental Methods.

Our gradostat flasks contained simple communities of the water flea *Daphnia pulex* consuming a mix of three algal species (*Pseudokirchneriella subcapitata, Scenedesmus quadricauda, Ankistrodesmus falcatus*). Configuration was controlled by connecting flasks in a linear configuration or a dendritic configuration. We also controlled functional connectivity contrasting metaecosystem dynamics when only nutrients moved versus the case when nutrients, resources and consumers moved. This experiment employed a 2×2×2 factorial design to test the importance of ecosystem trophic connectivity (a treatment considering movement of medium only vs. movement of media, phytoplankton and *Daphnia* between flasks) and metaecosystem configuration (linear or dendritic) on the stability of ecosystems with two levels of enriched medium input (regular and phosphorus-enriched). Each metaecosystem consisted of four “nodes” of 500 mL Erlenmeyer flasks with a foam stopper to allow for gas exchange, seeded initially with 100 mL algal mix (total average algal density of 2.22 x10^6^ +/- 1.3×10^4^ cells/mL) to which 50 adult *Daphnia* were added before topping off the flask to 500 mL with FLAMES media (21). Flasks were then connected by Tygon tubing and from an inflow reservoir of FLAMES medium (10 μgP/L) or enriched P (70 μgP /L) medium was pumped through the array of flasks using peristaltic pumps (Watson-Marlow 503S/RL and Rainin Dynamix RP-1). The dilution rate was 0.1 day^−1^ for all flasks in the linear configurations and the “hub” and “terminal” nodes of the dendritic configurations, and 0.05 day^−1^ for the “upstream” nodes in the dendritic configurations (Figure 1). To block the flow of organisms in the nutrient only connectivity treatment, outflow tubing was placed inside an 80-μm nylon mesh held in place with the stopper. Due to colony formation of the phytoplankton and clogging of the mesh this proved to be an effective retention mechanism also for the algal resources. In the trophic connectivity treatment, *D. pulex* were manually moved using a 2mL transfer pipette at a rate of ≈ 0.1 day^−1^ (20% were moved after each sampling count as sampling was only done every two days) in all linear nodes and the hub and terminal dendritic nodes, and ≈ 5% per day (10% moved after sampling) in the upstream dendritic nodes. Inflow stock solutions were prepared using FLAMES media (10 μgP/L). To increase P in the additionally nutrient-enriched treatment without changing pH, 132 μg/L of H_2_KPO_4_ and 168 μg/L HK_2_PO_4_ were added to our increased P treatment inflow stock solution. For the less-phosphorus treatment, no additional phosphorus was added, but 218 μg/L KCl were added to control for the K added to the high-P medium.

**Figure 1:**
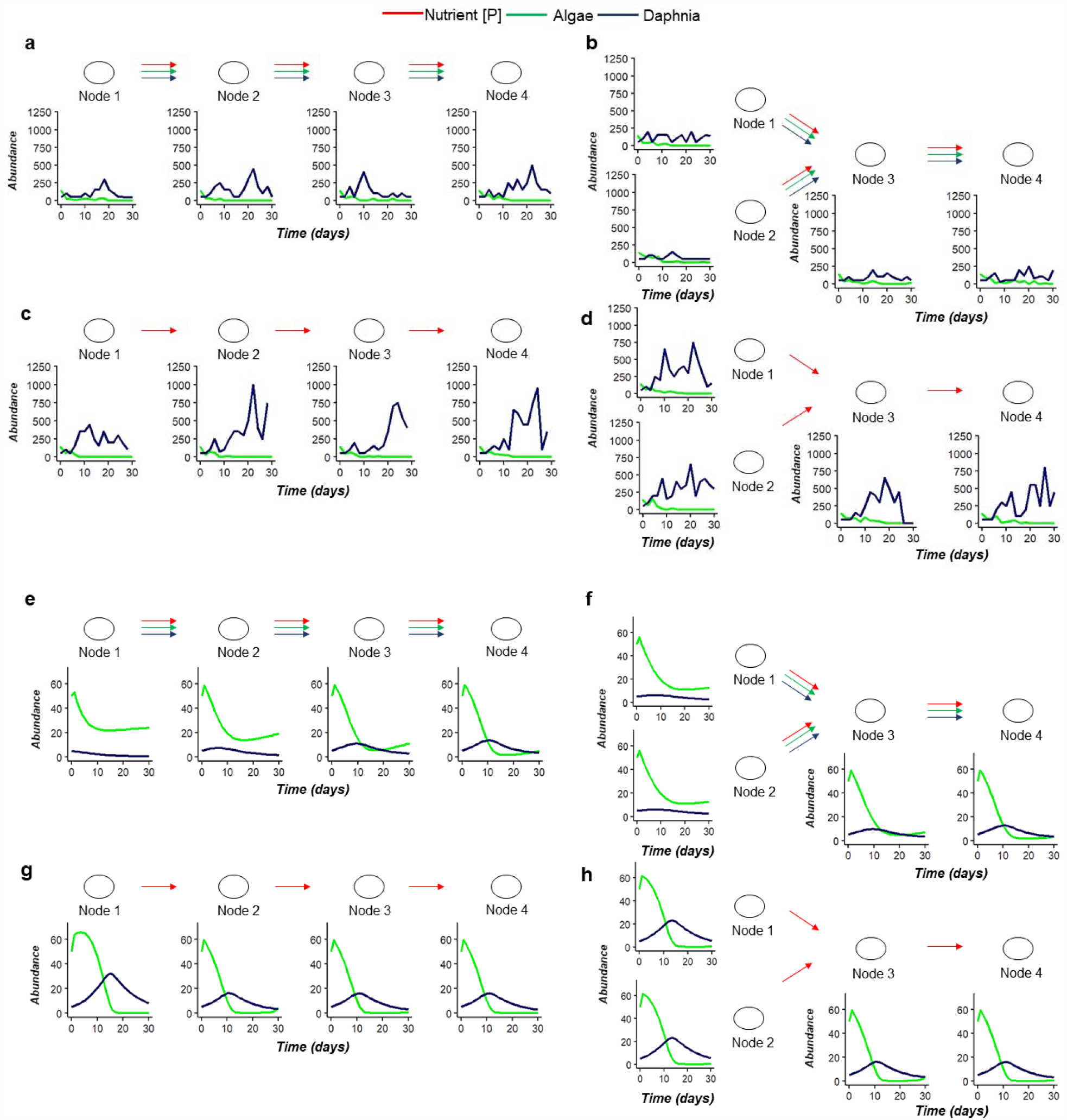
Population dynamics observed for Daphnia pulex (dark blue) and total algae concentration (green) within-node across metaecosystems by connectivity and configuration treatments based on an example experimental replicate (a-d) and from model simulations (e-h). For experimental data, D. pulex abundance is the estimated total number of animals in the flask based on our 50mL sample. Total algal abundance is the estimated number of cells/μL to fit the axis, based on a haemocytometer count. For model outputs, units are based on nutrient units with initial values of 50 for algae and 5 for Daphnia pulex.

### Experimental Sampling

The gradostats were sampled every other day for 30 days. In each node, the concentration of each algal species was measured using a haemocytometer. To estimate *Daphnia* population size, a 2mL plastic transfer pipette was used to gently agitate, then sample each node. The number of individuals and two age classes (adult or juvenile) in the pipette were determined and then replaced to the experimental flask. This process was repeated five times, and the total *D. pulex* count of the five samples was used to estimate *Daphnia* density/10mL (total number estimated per flask = total sampled count *50). A pilot testing this method proved it had an average error of 17.41 %. On Day 30 of the experiment, 40mL samples were taken from each flask to be analysed for total phosphorus concentration (TP). Phosphorus samples were analysed using a standard protocol (22) at the GRIL-Université du Québec à Montréal analytical laboratory.

### Statistical Analysis

To quantify the instability of *Daphnia* populations in experimental gradostats, we determined the peak total population size (as estimated by our 10mL samples) and the coefficient of variation (CV) of population size over the course of the experiment. These variables were calculated for each node within each gradostat, as well as in aggregate summed across all nodes. Similarly, CV and peak density were calculated for each species of alga but we report here values based on total algal density (sum of all species present).

All analyses of experimental gradostat data were conducted in R version 4 (23). For tests of statistical significance α was set to p = 0.05. To determine whether metaecosystem connectivity and configuration influenced local instability downstream of the nutrient enrichment source, we analysed the effects of our three experimental treatments and node position (1 upstream to 4 terminal) on *Daphnia* population peak and CV using generalized linear mixed effects models with block as a random factor and different error distributions depending on the outcome variable (Poisson distribution for *Daphnia* peak, with daphnia CV and algal CV log-transformed). We also tested for interactions between node position and each experimental treatment *(DV∼Node*(Connectivity+Configuration+Phosphorus)+(1*|*Block))*. Non-significant interactions were removed for final models

Final TP concentrations within each flask were compared among experimental treatments and node positions using a generalized linear mixed effects model with each treatment and node as fixed variables and block as a random effect *(TP∼Node* (Connectivity+Configuration+Phosphorus) +(1*|*Block))*

Non-significant interaction terms were removed for final statistical models.

We also measured instability at the scale of the entire gradostat. At the metaecosystem level, the effects of connectivity, configuration and phosphorus input treatments and their interactions on *Daphnia* peak and CV were tested using generalized linear mixed-effects models (package *lmer*). Block was included as a random factor in all models to account for any differences in initial algal concentrations or other temporal variability among blocks (*DV∼Connectivity*Configuration*Phosphorus +(1*|*Block)*. Different error distributions were applied depending on the response variable (Poisson distribution for peak metapopulation size, and log-transformation for CV), and non-significant interaction terms were removed one by one from the full statistical model. Goodness of fit of the final models were assessed with diagnostic plots of residuals.

### Metaecosystem Model

We examined the effects of trophic connectivity (i.e. how many trophic levels are moving from one node to the next) using a generic model to qualitatively assess patterns of ecosystem dynamics. We derived a model of 12 ordinary differential equations (ODEs) to describe a nutrient (*N*) - algae (*A*) - *Daphnia* (*D*) food chain repeated in four coupled flasks (subscripts 1, 2, 3, 4) with nutrient (only nutrients move) and trophic (nutrients, algae and *Daphnia* move) connectivity. The other treatment compared linear system with identical flowrates or dendritic systems with heterogenous flowrate configurations. This distinction of having either only nutrients move, or additionally *Daphnia* and algae move is akin to what is observed in aquatic ecosystems where organisms at different trophic levels may move at different rates and scales within river networks.

#### Linear Metaecosystems

We assume all flasks/“nodes” are homogenous in their initial *N, A* and *D* conditions. For linear configurations, inflow of medium (with nutrient concentration *N*_*sub*_) is into gradostat 1 at rate *δ*. In case of nutrient connectivity *δ*_*h*_ = 0, and only medium flows at rate *δ* from nodes 1 to 2, 2 to 3, 3 to 4, and out of 4. This modification of outflow from the terminal node was necessary due to the volume constraints of gradostat flasks. In case of trophic connectivity, nutrients, algae and *Daphnia* flow in the same manner at rate *δ* _*h*_ = *δ*. Assuming Monod-type uptake functions for both nutrients (by algae) and algae (by *Daphnia*) the model takes the form:

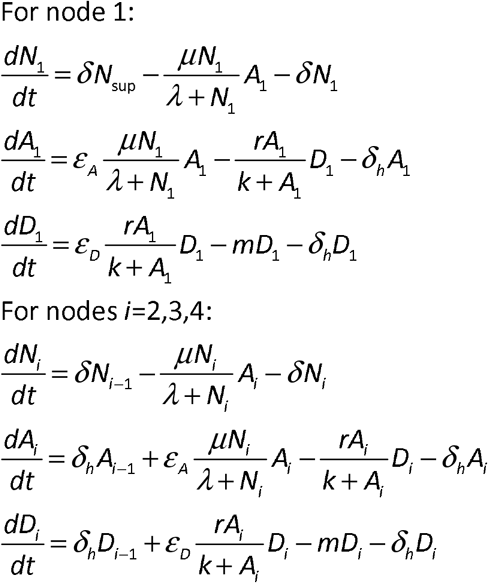

#### Dendritic Metaecosystem

We can also describe a dendritic gradostat model with heterogenous flow rates. In this configuration, flasks 1 and 2 are both recipients of the inflow medium, but at a slower rate *δ* ′ = *δ* /2 than in the linear configuration, and both flow out into flask 3, which then flows to flask 4 and out at the same rate as in the linear network. In analogy to the linear configuration, in case of trophic connectivity 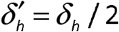, otherwise 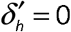. This modification of flow configuration in addition to rate in some nodes was necessary again due to the volume constraints of the gradostat flasks.

The **dendritic** model takes the form:

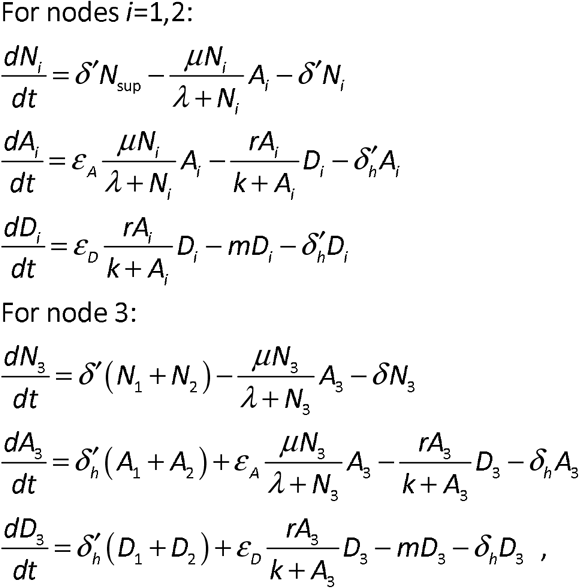

Initial state variable values were set to *N*_0_ = 25; *A*_0_ = 50; *D*_0_ = 5; in all four nodes of the gradostat system. These generic values in nutrient units are within the realm of possibility for a gradostat system but are not intended to quantitatively predict results or our gradostat experiments. Parameter estimates can be found in Table 1.

**Table 1:**
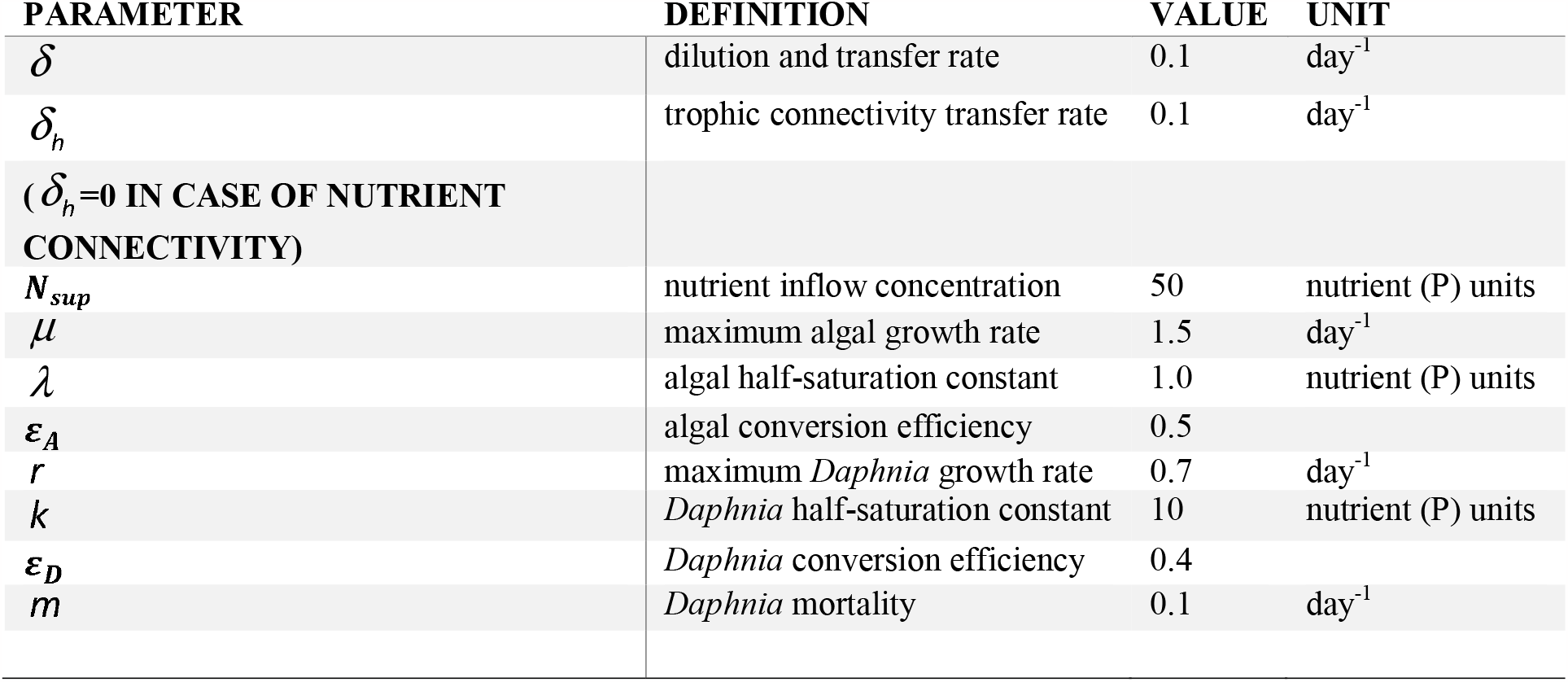
Parameter Estimates for Linear Configuration

The system of ODEs for the model was solved with ode45 in Matlab (R2022b) and model outputs for 30 days were obtained.

## Results

All reported values are means ± standard error unless otherwise stated.

### Phosphorus Gradients in Experimental Gradostats

Lack of sample preservation meant we were unable to obtain final TP concentration measurements for all our experimental nodes. With the samples we did obtain (one full replicate of each type of metaecosystem), we were unable to detect a significant difference in final P concentration among our high P and low P gradostats, indicating that all experienced a similar degree of disturbance in the form of nutrient enrichment, similar to the assumption of high nutrient loading in McCann et al (7). Nutrient-level was therefore removed from our statistical models for population dynamics and all P treatments were considered as enriched replicates. We detected a significant difference in P concentration by node position (1-4), regardless of metaecosystem connectivity or configuration treatment, with final TP increasing towards the terminal node (slope estimate=0.14±14.46, p=0.001), indicating that a nutrient gradient was established in our metaecosystems (Figure S1)

### Comparison Among Experimental Gradostat Nodes

In most of our gradostats, algae were grazed to undetectable levels before the end of the experiment. For *Daphnia* peak population size, we observed significant interactions between node position and connectivity (slope estimate = -0.11±0.007, p<0.001) and between node position and configuration (slope estimate = 0.06±.006, p<0.001). In general, peak population size was higher downstream (main effect slope estimate = 0.17±0.01), but this effect was stronger in gradostats with trophic connectivity than systems in which only nutrients were moved. This effect was also stronger in linear configurations compared to dendritic ones. Additionally, main effects of connectivity and configuration were detected, where population peaks were generally higher in nodes where only nutrients were moving (main effect slope estimate=1.09±0.02, p<0.001) and higher in linear compared to dendritic configurations (main effect slope estimate=0.09±0.02, p<0.001) (Figure 2a). No difference among any treatments, node position or interaction effects were detected for algal population sizes.

**Figure 2:**
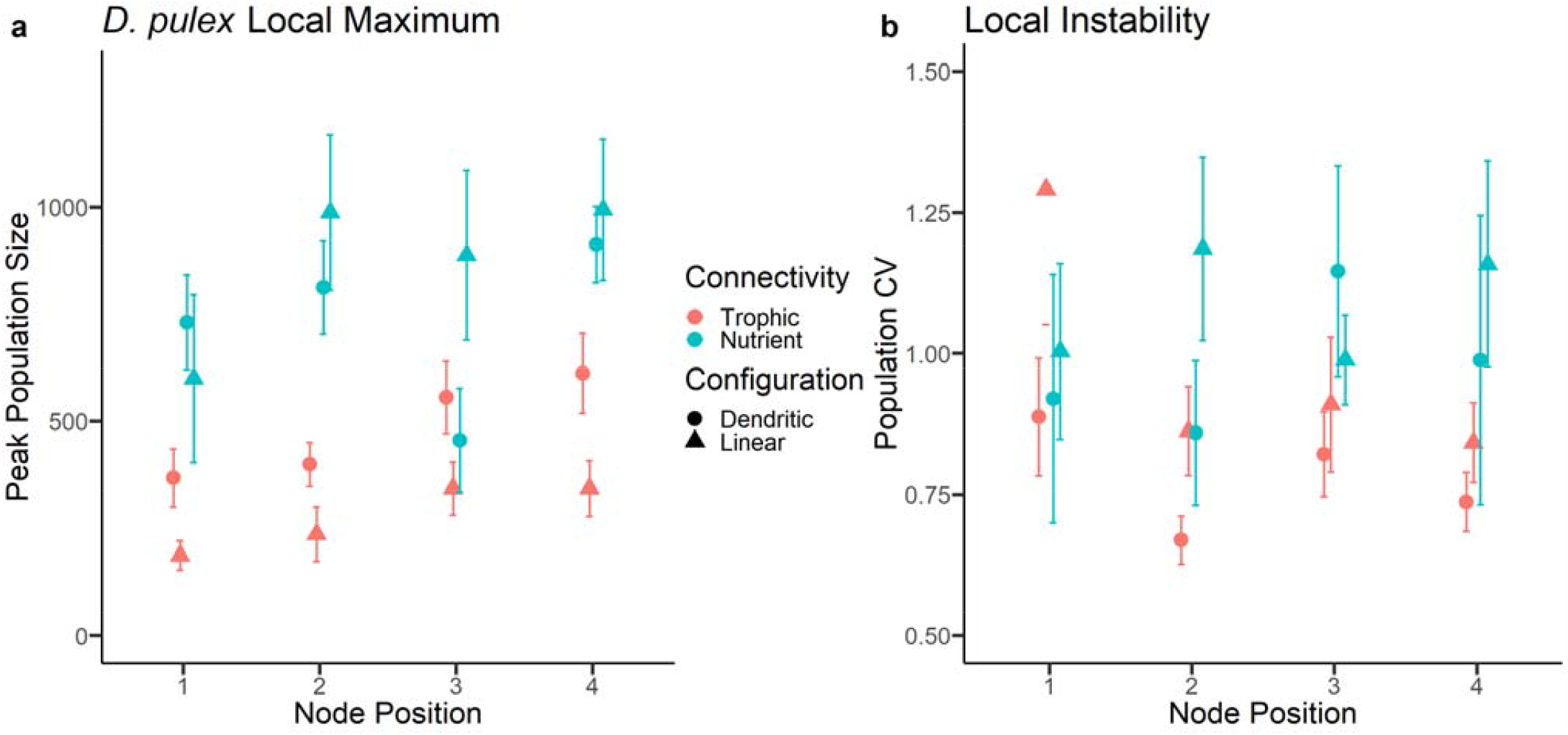
Within-node population instability (peak, a and cv, b) compared by metaecosystem connectivity and configuration. In general, Daphnia pulex population peak increases downstream from the source, but this effect is stronger in systems with trophic connectivity than those with nutrient connectivity and linear compared to dendritic networks. Population peaks are overall higher in networks with nutrient connectivity compared to those with trophic connectivity. For CV, instability increases with distance from nutrient input in nutrient connected systems, and decreases with distance from the source with trophically connected systems. CV is overall higher in nodes of systems with nutrient connectivity compared to those with trophic connectivity. Values are means ± 1 standard error, points are jittered horizontally to distinguish overlapping y-values.

For the CV of *Daphnia* populations, a significant interaction between connectivity and node position was detected (estimate=0.1±0.05 p=0.04), where instability increased far from the source of nutrient input in gradostats where only nutrients were moved, but decreased with distance from the source in gradostats where algae and *Daphnia* moved (Figure 2b). No difference among any treatments, node position or interaction effects were detected for algal CV.

### Comparison Among Experimental Networks

We also explored instability at the metaecosytem level. Similar to our results at the node level, a significant interaction of connectivity and configuration (estimate=0.96±0.02, p<0.001) on *Daphnia* peak was detected. Peak population size was higher in gradostats where only nutrients were moved than in gradostats where algae and *Daphnia* were moved, and this effect was more pronounced in linear systems compared to dendritic ones. A similar interaction of configuration and connectivity on metaecosystem *Daphnia* CV was also detected, where *Daphnia* CV was also greater for linear systems (0.66±0.34) compared to dendritic ones (0.55±0.03), but this effect was most pronounced in systems with nutrient connectivity (estimate=0.26±0.16, p<0.0001; Figure 3). No effect of any treatments or interactions was found for algal CV.

**Figure 3:**
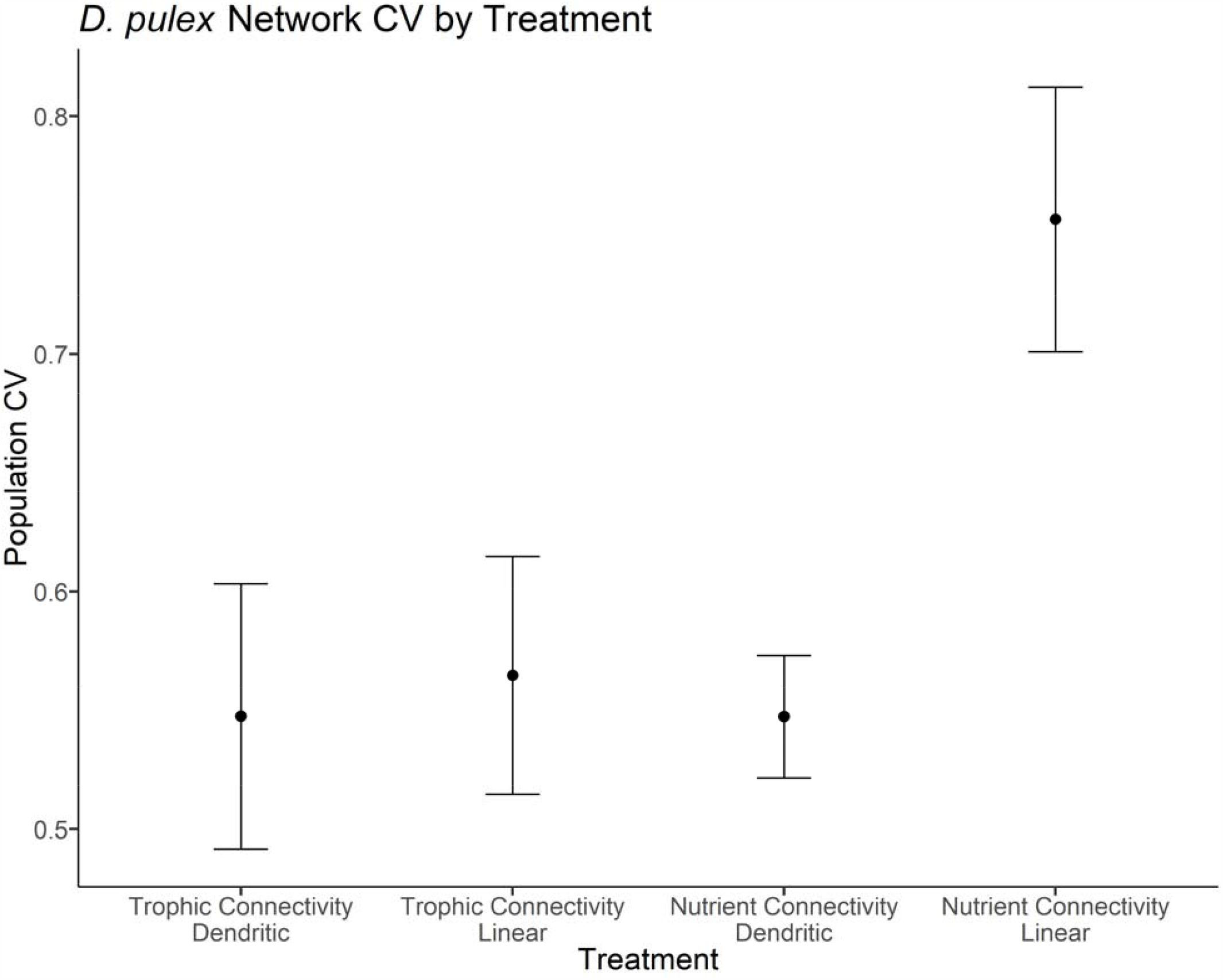
Interaction of connectivity and configuration on overall network instability (measured by Daphnia pulex population CV. Values are means ± 1 standard error. Nutrient connected networks were significantly less stable than networks with trophic connectivity where consumer and producers moved alongside nutrients, however metaecosystem configuration modified this relationship, with dendritic systems being stabilizing in nutrient connected systems compared to linear ones. For population peak (data not shown) a similar interaction was detected.

### Community Dynamics in Model Simulation

In general, our model simulations followed a similar pattern to our experimental results (Figure 1). In our mathematical model, *Daphnia* populations generally began in a growth phase in all nodes, reaching a population peak and then declining and stabilising, or going extinct. We did not predict or observe repeated oscillations of either species within the timeframe of our experiment (30 days). Upstream nodes in metaecosystems with trophic connectivity tended to remain relatively stable or decline. Overall, our model predictions indicated that *Daphnia* population size is amplified downstream in gradostats with trophic connectivity where consumers move with resources and nutrients, whereas this pattern is inverted in systems where only nutrients move, with consumer populations being largest in the upstream node. However, population sizes are higher overall in systems where only nutrients move. Heterogeneity in terms of slower upstream flow rates and a dendritic configuration dampens these effects such that peak population sizes are lower in downstream nodes of dendritic configurations compared to their linear counterparts.

At the metaecosystem level, our model predicted lower *Daphnia* population sizes in systems with trophic connectivity compared to those with only nutrients flowing between nodes.

## Discussion

We combined an experimental test with gradostats and a mathematical model to show that the stability of consumer-resource dynamics (*Daphnia* – algae dynamics) is influenced by the type of functional connectivity among communities and the configuration of nodes in the network. We compared our findings with simulations from a consumer-resource model of our metaecosystem. We hypothesised based on earlier theoretical work (7), that nutrient enrichment would drive local population instability in the “terminal” nodes of the network and that consumer movement would lead to greater network instability by synchronising consumer-resource dynamics within nodes. We also expected that spatial configuration of the network would modify these effects. Both our experimental data and model predictions demonstrate nutrient accumulation downstream from the enrichment source in connected ecosystems, with concentrations of P in terminal nodes almost four times that of the inflow, which may create the conditions for nutrient driven instability at locations far from the source (7). In keeping with this, we found that *Daphnia* instability was greater downstream, with peaks and CV of biomass up to twice those upstream indicating that this amplification of enrichment may cause bottom-up effects on community instability similar to those described in the paradox of enrichment (14). However, we found that connectivity with consumers moving with resources and nutrients stabilises community dynamics. Heterogenous flow rates among nodes provide a similar stabilising effect with dendritic configurations being more stable than linear ones, particularly for nutrient-connected systems.

### Connectivity and Configuration Significantly Impact Downstream Stability

We had hypothesised that trophic connectivity (*Daphnia* and algae moving with nutrients) would destabilise the dynamics by generating synchrony among nodes (13), because previous work predicted that both nutrient and consumer movement were capable of propagating instability through enriched systems. However, we found that nutrient connected gradostats actually had higher CV and population peaks that were almost twice the size of systems where consumers and algae moved. This instability was evident at the node and metaecosystem level. A key difference between our experiment and previous theoretical work (7) is that this earlier work considered consumer (i.e., *Daphnia*) movement alone, rather than with nutrient movement; this involves a decoupling of consumer density from nutrient concentration. Previous models by Gounand et al (24) predict that intermediate rates of consumer movement may have a stabilising effect due to negative density dependence, leading to smaller amplitude oscillations of consumer populations like those observed in our experiment. It is possible that within this experiment the immigration and emigration of consumers, which is known to stabilise population dynamics (25), meant that they were able to track and control algae resource growth.

We expected a dendritic configuration with heterogenous flowrates to “slow” the spread of instability throughout a network compared to a linear configuration. We see this effect in our results for peak *Daphnia* population size for the treatments in which only nutrients flowed. A dendritic configuration could have a more stabilising effect by increasing asynchrony among nodes because input is coming from multiple heterogeneous sources rather than a single source (7). In general, we saw that that nodes in dendritic systems were more stable than their linear counterparts, however the impacts were not as strong as those of functional connectivity treatments. Moreover, at the metaecosystem level, an interaction between connectivity and configuration was detected, where the stabilising effects of a dendritic configuration was stronger in linear nutrient connected systems compared to trophically connected systems. These findings therefore suggest that not only the degree of connectivity (in terms of movement of nutrients, resources, consumers, etc.), but the configuration of a metaecosystem in terms of direction and number of connections can affect stability, in line with theory (26).

### Model Simulations Predict Similar Patterns

Our confidence in this understanding of the experimental results is strengthened by the analysis of our model that shows that movement of consumers and algae with nutrients may have a stabilising effect on enriched ecosystems. Population sizes were lower in trophically connected systems, which may increase risk of extinction in a stochastic model. Although very few nodes went extinct in our experiment over 30 days, running our model for a longer period of time (Figure S2) suggests that trophic connectivity may lead to extinction within 100 days, while nutrient connected systems may lead to stable limit cycles that persist, despite the larger oscillations, at least in upstream nodes. Moreover, in our trophically connected simulations without outflow, our terminal nodes with large oscillations are the only ones to stabilise in the long-term (Figure S2). Running microcosms for a longer period of time might demonstrate the longer-term expectation of such a high degree of consumer removal. An analysis of longer-term dynamics also hints at different mechanisms for consumer instability under different connectivity regimes. In systems where only nutrients move, algae were first driven to large oscillations and very low densities, as predicted by the paradox of enrichment, whereas in systems where consumers move, algae eventually stabilised and consumers attained low densities, suggesting that connectivity may induce either top-down or bottom-up effects on ecosystem stability, depending on which trophic level(s) are moving.

Our model also demonstrated that node configuration may impact stability, with all dendritic systems having, smaller more stable populations than linear configurations. This effect was especially pronounced in trophically connected systems. The heterogenous flow rates dampened the effects of enrichment on downstream ecosystem stability compared to linear systems with homogenous flow rates. In these cases, the upstream nodes absorb most of the instability compared to downstream ones, with less amplification of instability downstream compared to linear systems; this results is in line with predictions made by McCann et al (7). However, as in our experimental results, even in dendritic systems connectivity had a large impact on local stability, with nutrient connectivity leading to greater instability than trophic connectivity. This implies that heterogeneity in flow rates has a minimal impact on metaecosystem instability compared to connectivity.

### Implications and Avenues for Future Work

Runoff from agricultural fertilizers and herbicides is a major contributor to nutrient enrichment in aquatic ecosystems (27, 28), as are point-source influxes from urban habitats (27, 29). We demonstrated experimentally that the effects of this enrichment on ecosystems may be propagated or even amplified at a distance far from the source, especially when the metaecosystem is connected by the movement of nutrients/water alone. These results are consistent with recent theory and imply that in some cases managing connectivity among ecosystems may dampen the negative impacts of enrichment on stability, in line with metapopulation theory (7).

We recognize a few key limitations of this work. First, our gradostats configurations are constrained in their equal for homogenous flow rates in/out of flasks, and future work could benefit from further examining how spatial heterogeneity in flow rates may influence overall metaecosystem stability. Furthermore, our model and experiment consider only one source point of nutrient enrichment, and real-world aquatic ecosystems may have many, which could also increase the accumulation of nutrients and therefore instability at the terminal node. Additionally, our model and experiment are simple in that they involve only two trophic levels and four nodes, with all nodes experiencing identical initial conditions. Investigating these dynamics in larger (both in size and number of nodes), more complex communities with multiple consumer species and at longer time scales would therefore be an important avenue for future work.

### Conclusion

Though theory has examined the impacts of consumer dispersal and nutrient enrichment on stability across metaecosytems (7, 24), this study is one of few (30) to empirically test this theory. We model and controlled an experimental system in which algae and consumers moved versus the cases where only medium moved with a flow of nutrients. We found that trophic connectivity (movement of algae and consumers) may stabilise an enriched system, but that metaecosystem configuration also may act to dampen instability. These findings have implications for agricultural and industrial practices, as well as how agriculture and urban systems are designed, as runoff and pollution are two large contributors of nutrient enrichment to aquatic ecosystems (7) and may also impact metaecosystem connectivity and configuration. We show that even a low level of nutrient input via fresh culture media is enough to destabilise ecosystems far from the input source, suggesting that even a moderate amount of runoff or inflow of nutrients similar to concentrations that are already present in the local environment could have similar effects. Additionally, we demonstrate that connectivity and configuration may diminish these effects, suggesting that preventing or slowing movement of some trophic levels at some nodes within a metaecosystem may dampen some of these impacts. Future work would benefit from scaling up these findings to larger-sized, more complex networks (node number and configuration) and communities (e.g. trophic structure). There is much more work to do to discover how anthropogenic inputs may destabilize metaecosystem dynamics across space and time via dispersal and community connectivity.

## Acknowledgements

We thank Libby Rothberg, Michelle Gros and Alessandra Loria for laboratory support. We thank Mark Romer and Mahnaz Mansoori for administration of the McGill University Phytotron. AG acknowledges the support of the Liber Ero Chair in Biodiversity Conservation. We thank Kevin McCann for stimulating discussions.

## Data Accessibility Statement

All raw data and codes necessary to reproduce our results will be uploaded publicly to Dryad digital repository upon acceptance of this manuscript.

## Tables and Figures

**Figure S1:**
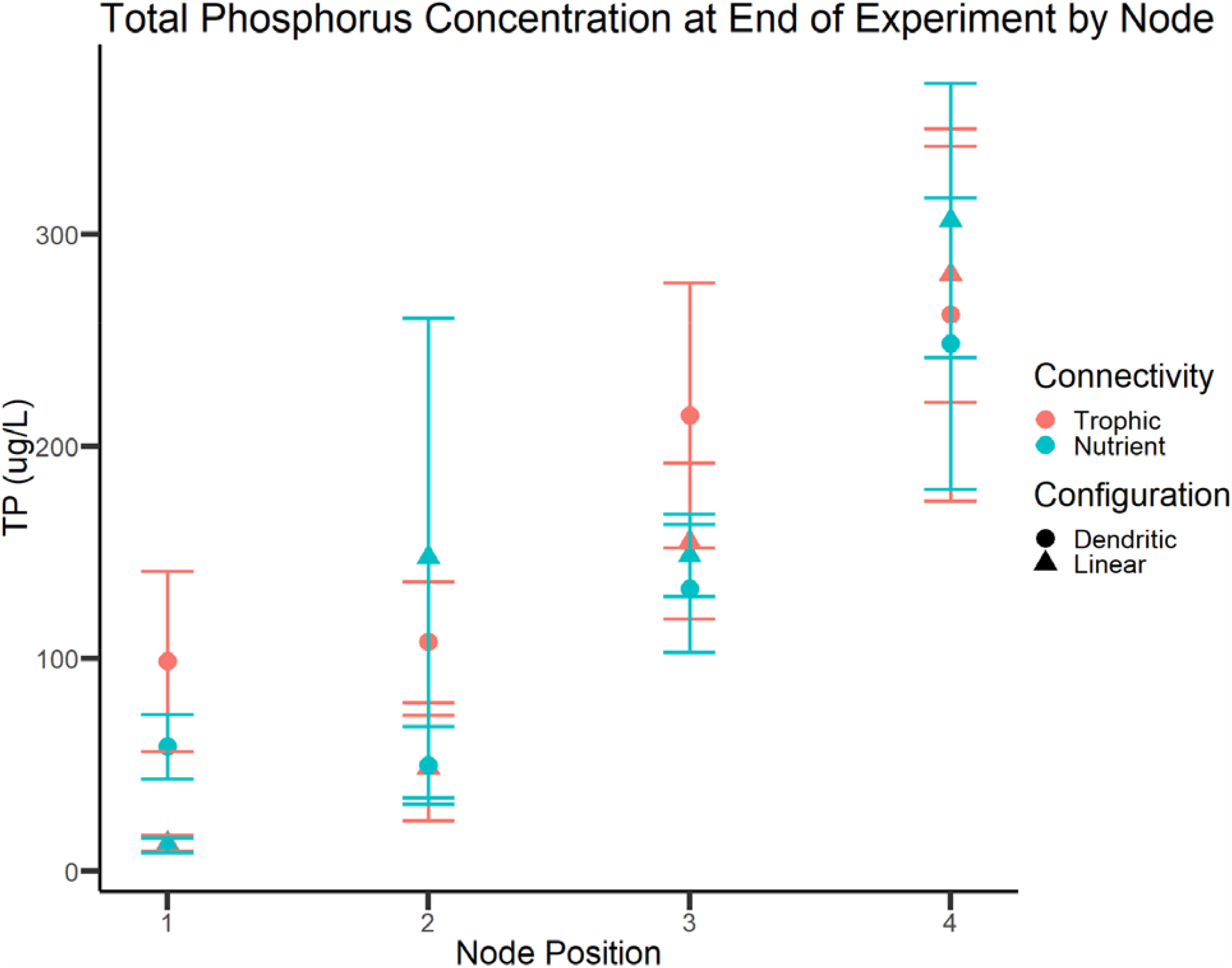
P gradient observed in all metaecosystems (final TP concentration mean ± 1 standard error) with total P accumulating in terminal node, with no difference among experimental treatments.

**Figure S2:**
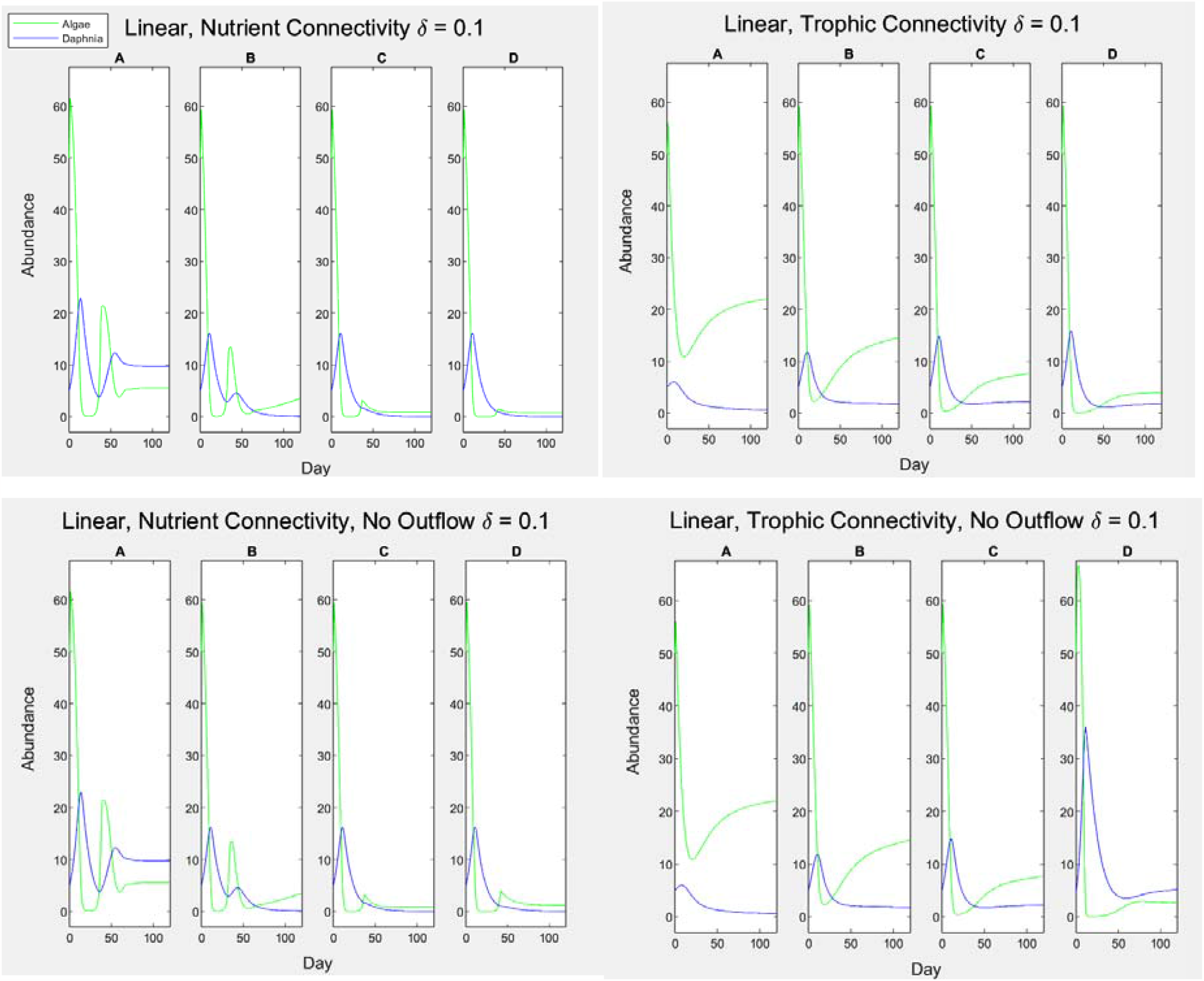
100-day (about the duration of a summer season in temperate zones) extension of the gradostat theoretical model, with and without outflow. Broad patterns are in line with our 30-model and experimental results. In both cases, nutrient connectivity leads to unstable oscillations which are propagated over the system. In trophically connected systems, instability is amplified further from the source, but overall lower than in nutrient connected systems.

## References

1. Ives AR, Carpenter SR. Stability and diversity of ecosystems. Science. 2007;317(5834):58–62.

2. McCann KS. The diversity–stability debate. Nature. 2000;405(6783):228–33.

3. Donohue I, Hillebrand H, Montoya JM, Petchey OL, Pimm SL, Fowler MS, et al. Navigating the complexity of ecological stability. Ecology Letters. 2016;19(9):1172–85.

4. Loreau M, Mouquet N, Holt RD. Meta-ecosystems: a theoretical framework for a spatial ecosystem ecology. Ecology Letters. 2003;6(8):673–9.

5. Thompson PL, Gonzalez A. Dispersal governs the reorganization of ecological networks under environmental change. Nature Ecology & Evolution. 2017;1(6):1–8.

6. Pedersen EJ, Marleau JN, Granados M, Moeller HV, Guichard F. Nonhierarchical dispersal promotes stability and resilience in a tritrophic metacommunity. The American Naturalist. 2016;187(5):E116–E28.

7. McCann KS, Cazelles K, MacDougall AS, Fussmann GF, Bieg C, Cristescu M, et al. Landscape modification and nutrient-driven instability at a distance. Ecology Letters. 2021;24(3):398–414.

8. Rooney N, McCann K, Gellner G, Moore JC. Structural asymmetry and the stability of diverse food webs. Nature. 2006;442(7100):265–9.

9. Bell G, Fugère V, Barrett R, Beisner B, Cristescu M, Fussmann G, et al. Trophic structure modulates community rescue following acidification. Proceedings of the Royal Society B. 2019;286(1904):20190856.

10. Fugère V, Hébert M-P, Da Costa NB, Xu CC, Barrett RD, Beisner BE, et al. Community rescue in experimental phytoplankton communities facing severe herbicide pollution. Nature Ecology & Evolution. 2020;4(4):578–88.

11. Firkowski CR, Thompson PL, Gonzalez A, Cadotte MW, Fortin MJ. MultiLJtrophic metacommunity interactions mediate asynchrony and stability in fluctuating environments. Ecological Monographs. 2022;92(1):e01484.

12. Quévreux P, Loreau M. Synchrony and Stability in Trophic Metacommunities: When Top Predators Navigate in a Heterogeneous World. Frontiers in Ecology and Evolution. 2022;10.

13. Gouhier TC, Guichard F, Gonzalez A. Synchrony and stability of food webs in metacommunities. The American Naturalist. 2010;175(2):E16–E34.

14. Rosenzweig ML. Paradox of enrichment: destabilization of exploitation ecosystems in ecological time. Science. 1971;171(3969):385–7.

15. Levin SA, Segel LA. Hypothesis for origin of planktonic patchiness. Nature. 1976;259(5545):659-.

16. Lovitt RW, Wimpenny JWT. The Gradostat: a Bidirectional Compound Chemostat and its Application in Microbiological Research. Microbiology. 1981;127(2):261–8.

17. Gaki A, Theodorou A, Vayenas DV, Pavlou S. Complex dynamics of microbial competition in the gradostat. Journal of Biotechnology. 2009;139(1):38–46.

18. Smith H, Waltman P. The gradostat: a model of competition along a nutrient gradient. Microbial Ecology. 1991;22(1):207–26.

19. Gülzow N, Wahlen Y, Hillebrand H. Metaecosystem Dynamics of Marine Phytoplankton Alters Resource Use Efficiency along Stoichiometric Gradients. The American Naturalist. 2019;193(1):35–50.

20. Smith HAL. Microbial growth in periodic gradostats. The Rocky Mountain Journal of Mathematics. 1990;20(4):1173–94.

21. Celis-Salgado MP, Cairns A, Kim N, Yan ND. The FLAMES medium: a new, soft-water culture and bioassay medium for Cladocera. SIL Proceedings, 1922-2010. 2008;30(2):265–71.

22. Wetzel RG, Likens GE. Limnological Analyses: Springer New York; 2013.

23. R Core Team. R: A language and environment for statistical computing. Vienna, Austria: R Foundation for Statistical Computing; 2020.

24. Gounand I, Mouquet N, Canard E, Guichard F, xe, xe, et al. The Paradox of Enrichment in Metaecosystems. The American Naturalist. 2014;184(6):752–63.

25. Gonzalez A, Mouquet N, Loreau M. Biodiversity as spatial insurance: the effects of habitat fragmentation and dispersal on ecosystem functioning. 2009.

26. Gilarranz LJ, Rayfield B, Liñán-Cembrano G, Bascompte J, Gonzalez A. Effects of network modularity on the spread of perturbation impact in experimental metapopulations. Science. 2017;357(6347):199–201.

27. Bashir I, Lone FA, Bhat RA, Mir SA, Dar ZA, Dar SA. Concerns and threats of contamination on aquatic ecosystems. Bioremediation and biotechnology: sustainable approaches to pollution degradation. 2020:1–26.

28. Pericherla S, Karnena MK, Vara S. A review on impacts of agricultural runoff on freshwater resources. Int J Emerg Technol. 2020;11:829–33.

29. Wurtsbaugh WA, Paerl HW, Dodds WK. Nutrients, eutrophication and harmful algal blooms along the freshwater to marine continuum. WIREs Water. 2019;6(5):e1373.

30. Laan E, Fox JW. An experimental test of the effects of dispersal and the paradox of enrichment on metapopulation persistence. Oikos. 2020;129(1):49–58.

